# Oxamniquine Derivatives Overcome Praziquantel Treatment Limitations for Schistosomiasis

**DOI:** 10.1101/2023.05.23.541778

**Authors:** Sevan N. Alwan, Alexander B. Taylor, Jayce Rhodes, Michael Tidwell, Stanton F. McHardy, Philip T. LoVerde

**Affiliations:** Departments of Biochemistry and Structural Biology, University of Texas Health at San Antonio; San Antonio, Texas 78229, USA; Biology Core Facilities, University of Texas Health at San Antonio; San Antonio, Texas 78229, USA; Center for Innovative Drug Discovery, Department of Chemistry, University of Texas at San Antonio; San Antonio, Texas 78249, USA

**Keywords:** *Schistosoma*, drug discovery, crystallography, drug resistance, RNA interference (RNAi)

## Abstract

Human schistosomiasis is a neglected tropical disease caused by *Schistosoma mansoni, S. haematobium,* and *S. japonicum.* Praziquantel (PZQ) is the method of choice for treatment. Due to constant selection pressure, there is an urgent need for new therapies for schistosomiasis. Previous treatment of *S. mansoni* included the use of oxamniquine (OXA), a drug that is activated by a schistosome sulfotransferase (SULT). Guided by data from X-ray crystallography and *Schistosoma* killing assays more than 350 OXA derivatives were designed, synthesized, and tested. We were able to identify CIDD-0150**610** and CIDD-0150**303** as potent derivatives *in vitro* that kill (100%) of all three *Schistosoma* species at a final concentration of 71.5 µM. We evaluated the efficacy of the best OXA derivates in an *in vivo* model after treatment with a single dose of 100 mg/kg by oral gavage. The highest rate of worm burden reduction was achieved by CIDD-150**303** (81.8%) against *S. mansoni*, CIDD-0149**830** (80.2%) against *S. haematobium* and CIDD-066**790** (86.7%) against *S. japonicum*. We have also evaluated the ability of the derivatives to kill immature stages since PZQ does not kill immature schistosomes. CIDD-0150**303** demonstrated (100%) killing for all life stages at a final concentration of 143 µM *in vitro* and effective reduction in worm burden *in vivo* against *S. mansoni*. To understand how OXA derivatives fit in the SULT binding pocket, X-ray crystal structures of CIDD-0150**303** and CIDD-0150**610** demonstrate that the SULT active site will accommodate further modifications to our most active compounds as we fine tune them to increase favorable pharmacokinetic properties. Treatment with a single dose of 100 mg/kg by oral gavage with co-dose of PZQ + CIDD-0150303 reduced the worm burden of PZQ resistant parasites in an animal model by 90.8%. Therefore, we conclude that CIDD-0150**303**, CIDD-0149**830** and CIDD-066**790** are novel drugs that overcome some of PZQ limitations, and CIDD-0150**303** can be used with PZQ in combination therapy.

**Author Summary:** Human schistosomiasis is a neglected tropical disease caused by parasitic worms in the genus *Schistosoma*. Human schistosomiasis is caused mainly by three major species: *S. mansoni, S. haematobium,* and *S. japonicum.* It affects some 229 million people in 78 countries. Currently, there is no effective vaccine against human schistosomiasis. Praziquantel is the method of choice for treatment and evidence for drug resistance has been reported. Our focus is drug discovery for schistosomiasis. Our project team is designing, synthesizing, and testing reengineered derivatives of oxamniquine against the three human species of *Schistosoma*. The aim is to develop a new drug for schistosomiasis to overcome developing resistance and improve efficacy. We developed and identified compounds that kill all three human *Schistosoma* species in addition to a PZQ-resistant strain in animal models. Additionally, animal studies demonstrate that combination treatment of reengineered oxamniquine drugs and praziquantel effectively reduced the infection with a praziquantel resistant strain in infected mice.

## Introduction

Human schistosomiasis is a neglected tropical disease caused by parasitic flatworms in the genus *Schistosoma*. Human schistosomiasis is caused mainly by three major species: *S. mansoni, S. haematobium,* and *S. japonicum*. It affects some 229 million people globally [1–3]. Of those infected 20,000-200,000 people are estimated to die from the disease annually [4–6]. However, the major impact of schistosomiasis is life years lost due to morbidity. The DALYs index (“Disability-Adjusted Life Years”) for schistosomiasis is estimated at 1.9 million [7]. *Schistosoma* has a complex life cycle that involves freshwater snails as intermediate hosts and humans among others as a final host. Three stages of *Schistosoma* life cycle live in an infected human host; eggs, juvenile worms, and adult worms. The infection leads to periportal fibrosis, portal hypertension, liver and spleen enlargement and the serious sequelae of esophageal and upper gastrointestinal varices, recurrent hematemesis, abdominal ascites and urogenital involvement such as, bladder deformity, hydronephrosis, hematuria, female genital schistosomiasis, infertility, increased risk of HIV-1 transmission, and squamous cell carcinoma of the bladder [8–11]. There are also systemic morbidities associated with *Schistosoma* infection such as anemia, growth stunting, impaired cognition, undernutrition, diarrhea, and decreased physical fitness [11]. Currently, there is no effective vaccine against human schistosomiasis. Only one treatment, praziquantel (PZQ) is available. Although PZQ is effective against all three species, the reported cure rates are 60-90% [12], PZQ is not effective against juvenile worms, it does not prevent reinfection, and evidence for drug resistance has been reported [13–17].

Oxamniquine (OXA) is a drug that was used previously to treat millions of people with *S. mansoni* with cure rates similar to PZQ [18, 19]. OXA is effective against adult stage schistosomes, and evidence of drug resistance against OXA in the laboratory and in the field has been demonstrated [20, 21]. A study by Cioli et al. demonstrated that OXA resistance is a double recessive trait. With this information Valentim et al. identified the gene responsible for OXA resistance [22, 23]. OXA is activated by *S. mansoni* sulfotransferase (*Sm*SULT) via transiently adding a sulfate to a hydroxy-methyl group. The activated form of OXA undergoes nucleophilic attack by macromolecules such as DNA, resulting in killing of *S. mansoni* [22, 24]. Alternatively, the sulfur group will decay and activated OXA acts as an electrophile forming adducts with macromolecules and interfering with schistosome metabolism [25]. Although sulfotransferase orthologs are expressed by *S. haematobium* and *S. japonicum,* OXA is not effective against these two species [24]. However, differences in sulfotransferase enzyme efficiency, variation in detoxification processes between species, and differences in sulfotransferase concentration remain possible explanations for species-specific resistance and may be interdependent in establishing OXA toxicity. Therefore, one answer to the question is that OXA kills *S. mansoni* but not *S. haematobium* or *S. japonicum* because it does not fit into the SULT binding pocket productively and does not get activated to a sufficiently toxic level [26].

Due to the danger of resistance to the monotherapy praziquantel, developing a novel drug will have a significant impact on global human health and will lead to improved treatments for *Schistosoma* to reduce the morbidity, mortality, and transmission rates associated with these devastating infections. An iterative process for drug development has been used to identify derivatives of OXA that demonstrate effective killing against all three human species of the parasite [25, 27, 28]. From these, we were able to identify CIDD-0066**790** and it’s (*R*)-enantiomer CIDD–0072**229**, both of which demonstrated broad species killing activity: *S. mansoni* (75%), *S. haematobium* (40%) and *S. japonicum* (83%) and *S. mansoni* (93%), *S. haematobium* 95% and *S. japonicum* 80%, respectively in an *in vitro* killing assay [27, 28]. Recently, we were able to identify the derivative CIDD-0149**830** that kills 100% of the *S. mansoni*, *S. haematobium* and *S. japonicum* worms *in vitro* within one week compared to 14 days for OXA to kill *S. mansoni* [28]. Our goal is to develop a novel therapeutic that will kill all three species of *Schistosoma* that has a mechanism of action different from PZQ to overcome the potential for resistance and enhance efficacy. In this paper, we present data identifying 2 OXA derivatives that kill all human species and work against liver stage schistosomes. Thus, have the potential to improve chemotherapy.

## Results

We previously identified CIDD-0149**830** (referred to as **830**) that shows 100% pan-specific killing activity *in vitro*. In this study we have identified an additional two derivatives that show 100% pan-specific killing activity after treatment for 45 minutes at a final concentration of 143 µM *in vitro* (Fig 1), CIDD-0150**610** (referred to as **610**) and CIDD-0150**303** (referred to as **303**) of which **303** is an enantiomer of **830** (S1 Table). **610** and **303** were able to kill 100% of *S. mansoni* at 143 µM within 24 hours in an *in vitro* killing assay.

**Fig 1:**
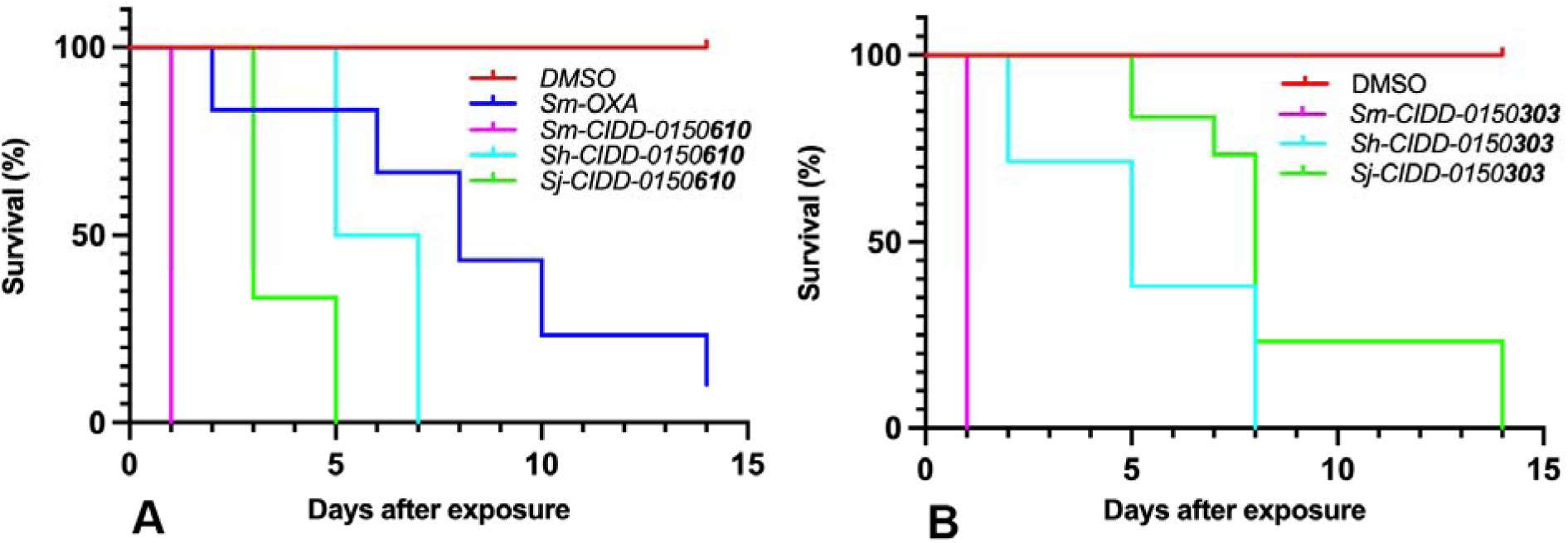
Kaplan-Meier Curves Demonstrate the Ability of OXA Derivatives to Kill Adult Schistosomes *In Vitro*. A. CIDD-0150**<u>610</u>**, B. CIDD-0150**<u>303</u>**. Both compounds kill 100% *of S. mansoni, S. haematobium* and *S. japonicum* compared to OXA that kills 90% of *S. mansoni in vitro*. OXA derivatives were tested against 10 adult male worms per well. All derivatives were solubilized in 100% DMSO, administered at a final concentration of 143 µM per well for 45 minutes, washed 3 timess with media. 45-minute exposure mimics the exposure time in a human (D. Cioli, pers commun.). All screens were performed in experimental and biological triplicate. Survival was plotted as a percentage over time using Prism/Curve Comparison/ Long-rank (Mantel-cox) test. The p-value threshold for each derivative compared to DMSO was <0.001.

In order to determine the minimum dose for those derivatives that demonstrate the best killing (100%) of the three species within 14 days of incubation, we evaluated a dose response using 143 µM, 71.5 µM, 35.75 µM and 14.3 µM to determine the best concentration for killing. Fig 2 shows the ability of **303** and **610** to kill 100% of *S. mansoni, S. haematobium,* and *S. japonicum* worms at 71.5 µM and about 50% at 35.75 µM (S1 Fig). Fig 2 also shows a final concentration of 71.5 µM of OXA was able to only kill 40% of *S. mansoni* within 14 days.

**Fig 2:**
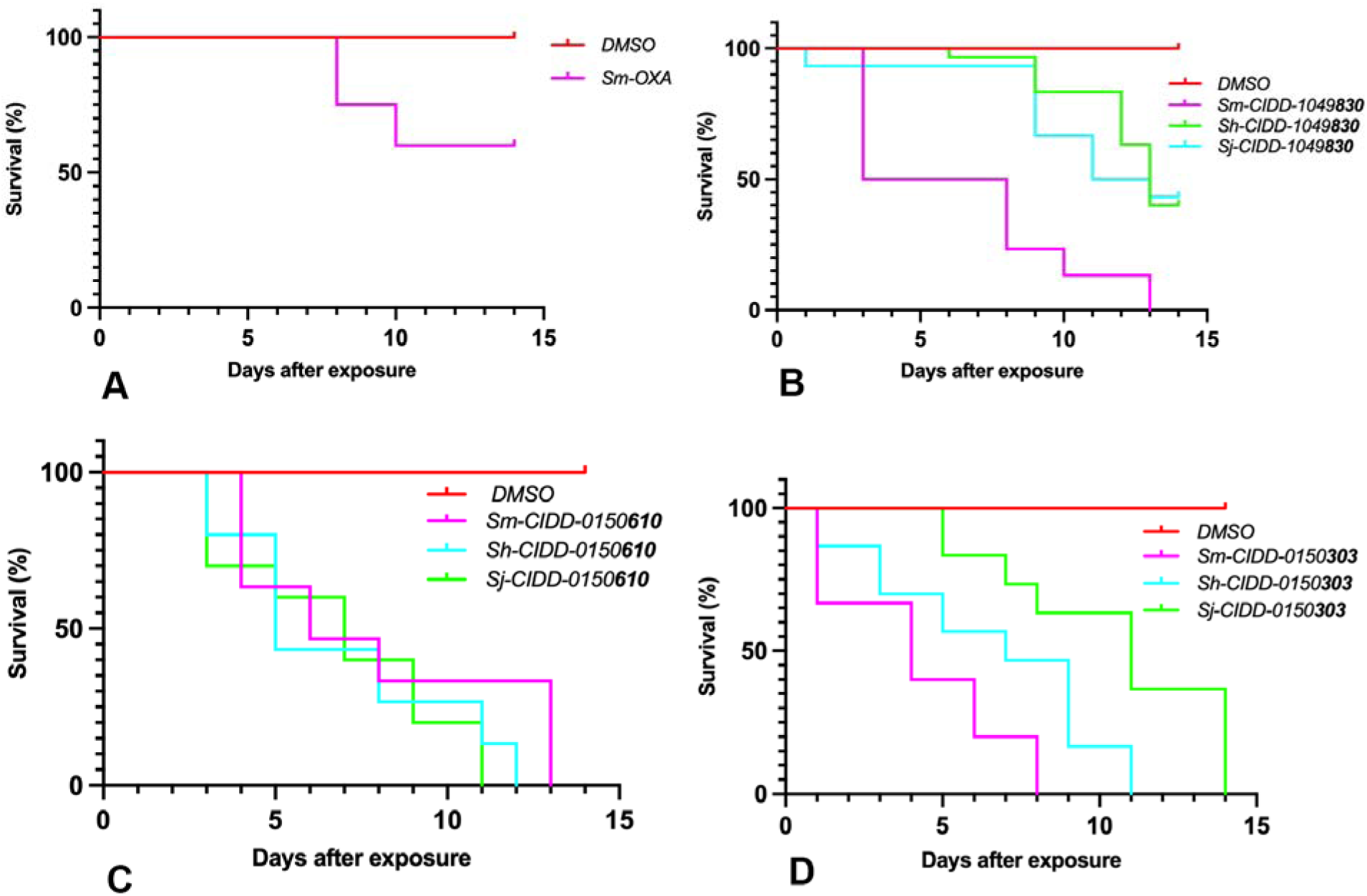
Kaplan-Meier Curves Demonstrate the Effect of Final Concentration Of 71.5 µm Per Well. A. OXA, B. CIDD-0149**<u>830</u>**, C. CIDD-0150**<u>610</u>**, and D. CIDD-0150**<u>303</u>**. **610** and **303** will kill 100% of *S. mansoni, S. haematobium* and *S. japonicum*. **830** will kill 100% of *S. mansoni,* 60% of *S. haematobium* and 56.7% of *S. japonicum*. OXA will kill 40% of *S. mansoni* OXA and OXA derivatives were tested against 10 adult male worms per well. All derivatives were solubilized in 100% DMSO. The worms were treated for 45 minutes, washed 3 times with media. 45-minute exposure mimics the exposure time in a human All screens were performed in experimental and biological triplicate. Survival was plotted as a percentage over time using Prism/Curve Comparison/ Long-rank (Mantel-cox) test. The p-value threshold for each derivative compared to DMSO was <0.001.

We have tested the ability of OXA and OXA derivatives: **830**, **610** and **303** to kill both schistosome genders. The drugs **610** and **303** kill 100% at a final concentration of 143 µM of unpaired female and male worms each from bisex infection and female and male worms in worm pairs. Interestingly, **303** was able to kill them all within 24 hours at 143 µM. Paired male worms were less susceptible to OXA (Fig 3).

**Fig 3:**
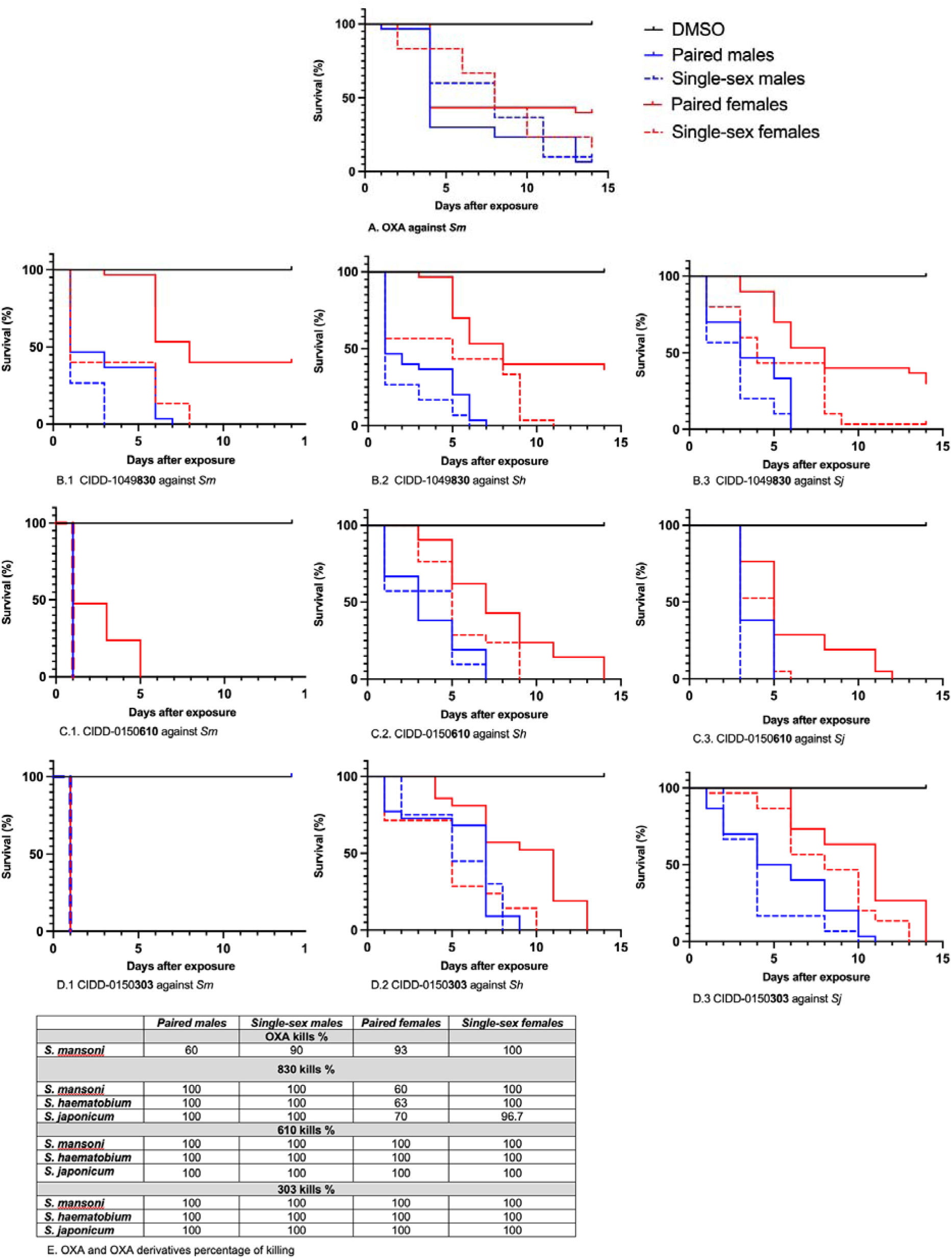
Kaplan-Meier Curves Demonstrate the Ability of OXA And OXA Derivatives to Kill Both Genders. A. OXA to kill both genders of *S. mansoni* B. CIDD-0149**<u>830</u>** against both genders of B.1. *S. mansoni*, B.2. *S. haematobium,* and B.3*. S. japonicum.* C. CIDD-0150**<u>610</u>** against both genders of C.1. *S. mansoni*, C.2. *S. haematobium,* and C.3*. S. japonicum.* D. CIDD-0150**<u>303</u>** against both genders of D.1. *S. mansoni*, D.2. *S. haematobium,* and D.3*. S. japonicum.* E. Percentages of worm killing. OXA and OXA derivatives were tested against 10 adults of single sex-female and male worms and female and male worms in worm pairs per well. **610** and **303** kill 100% of both gender from *S. mansoni, S. haematobium* and *S. japonicum*. **830** kills 100% of paired males, and single-sex females and 96.7% of single-sex females. Paired females were less susceptible to **830**. OXA demonstrates the expected level of killing against single-sex males and paired females from *S. mansoni*, the drug is effective 100% against single-sex females and less effective 60% against paired males. All derivatives were solubilized in 100% DMSO and administered at a final concentration of 143 µM per well for 45 minutes, washed 3 times with media. 45-minute exposure mimics the exposure time in a human. All screens were performed in experimental and biological triplicate. Survival was plotted as a percentage over time using Prism/Curve Comparison/ Long-rank (Mantel-cox) test. The p-value threshold for each derivative compared to DMSO was <0.00.

To test the efficacy of OXA derivates in an *in vivo* model, five mice per group were infected with 80 *S. mansoni* cercariae, treated by oral gavage with 100 mg/kg of OXA, **830**, **610**, and **303** at day 45 post-exposure. Ten days after treatment with **830** and **303** the worm burden was reduced by 72.3% *P=0.012* and 81.8% *P=0.0017,* respectively. The reduction in worm burden after **610** treatment was 47% *P=0.054* and was not significant. However, OXA reduced the number of worms by 93% *P= 0.0002* (Fig 4A). Five hamsters per group were infected with 100 cercariae of *S. haematobium*, treated 90 days later with 100 mg/kg of **830**, **610**, and **303**. Treatment with all compounds showed significant killing. The reduction in the number of collected worms after **830** treatment was 80.2% *P=0.0001*. Furthermore, **610** and **303** showed significant killing for *S. haematobium* infection 69.1% and 60%, respectively (Fig 4B). To evaluate OXA derivates against *S. japonicum* in infected hamsters, five animals per group were treated with 100 mg/kg of **830**, **610**, **303**, and CIDD-066**790** (referred to as **790**) at day 30 post-exposure. CIDD-066**790** is a derivative that was identified previously and demonstrated broad species killing activity. After *S. japonicum* worm collection, we obtained a reduction in worm burden of 38.3% *P= 0.00443* with **830**, 61% *P= 0.0019* with **610**, 31% *P= 0.121* with **303**, and 86.7 % *P= 0.0003* **790** compared to control animals (Fig 4C).

**Fig 4:**
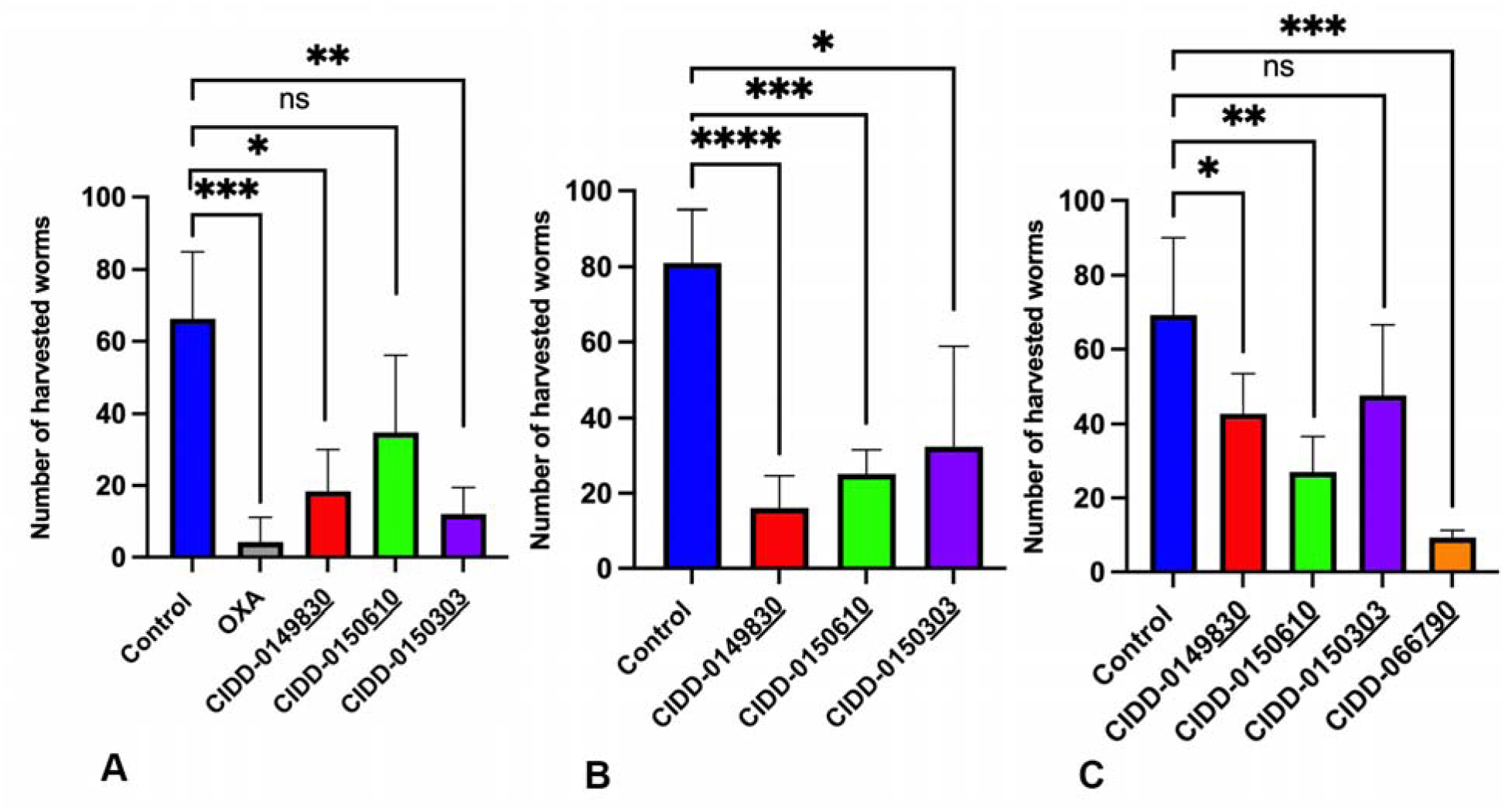
Effect of OXA Derivatives on A. *S. mansoni* B. *S haematobium* C*. S. japonicum* Infected Animals. Five mice per group were infected with *S. mansoni*, 5 hamsters per group were infected with *S. haematobium,* and 5 hamsters per group were infected *with S. japonicum* Worms were collected 10 days after treatment with a single dose of 100 mg/kg by oral gavage compared to the untreated control group. Prism/unpaired t test *(P<0.05)*.

One of the PZQ treatment limitations is that PZQ does not kill immature schistosomes and therefore, we have also focused on derivatives that will kill immature, liver stage schistosomes [29, 30]. Therefore, we treated liver stages with 143 µM of **830**, **610** and **303** in an *in vitro* assay. **303** treatments in an *in vitro* assay leads to 100% killing of liver stage 20-28 dpi worms in 2 days (Fig 5).

**Fig 5:**
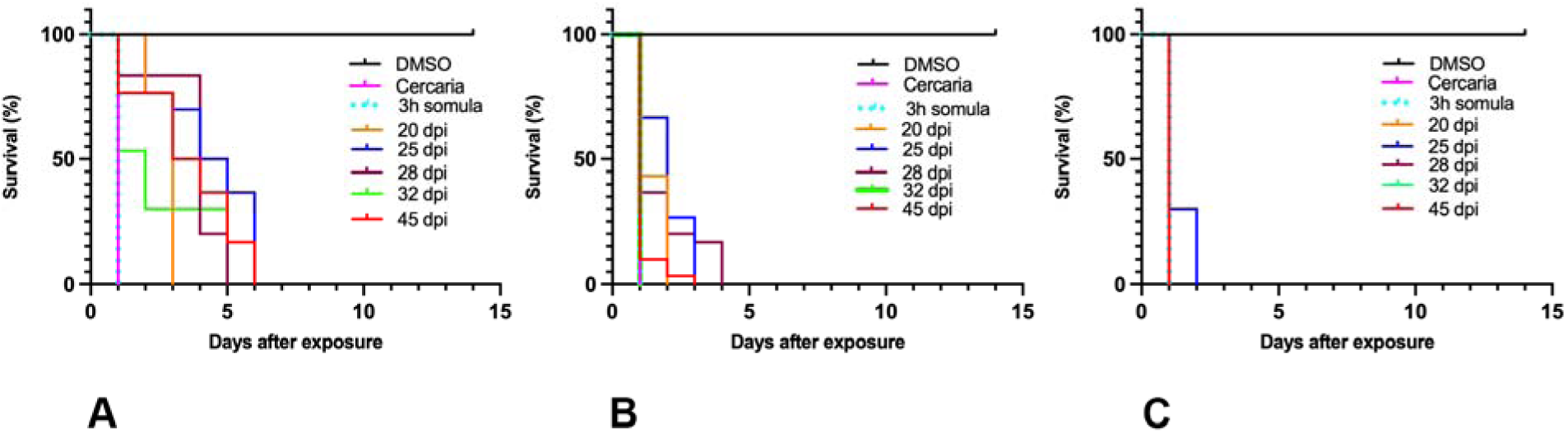
Kaplan-Meier Curves Demonstrate the Ability of OXA Derivatives to Kill All Life Stages of *S. mansoni*. A. CIDD-0149**<u>830</u>**, B. CIDD-0150**<u>610</u>**, and C. CIDD-0150**<u>303</u>** OXA derivatives were tested against cercaria, 3-hour schistosomula, and male worms for 20 dpi, 25 dpi, 28 dpi, 32 dpi and 45 dpi. OXA derivatives demonstrate 100% of killing for all life stages within 2-6 days. All derivatives were solubilized in 100% DMSO and administered at a final concentration of 143 µM per well for 45 minutes, washed 3 times with media. 45-minute exposure mimics the exposure time in a human. Each well contained 10 male adult worms. All screens were performed in experimental and biological triplicate. Survival was plotted as a percentage over time using Prism/Curve Comparison/ Long-rank (Mantel-cox) test. The p-value threshold for each derivative compared to DMSO was <0.001.

We tested the ability of **303** to kill juvenile worms in an *in vivo* study. Five mice per group were infected with 80 *S. mansoni* cercariae, treated with **303** at 100 mg/kg as a single oral dose on 20 dpi, 25 dpi, 28 dpi and 32 dpi and perfused 45 dpi. **303** reduced the worm burden significantly on 25 dpi by 63.8% *P= 0.0001*, 28 dpi by 48.9% *P= 0.000,* and 32 dpi by 54.1% *P= 0.0005* (Fig 6). At 20 dpi, worm burden reduction was not significant.

**Fig 6:**
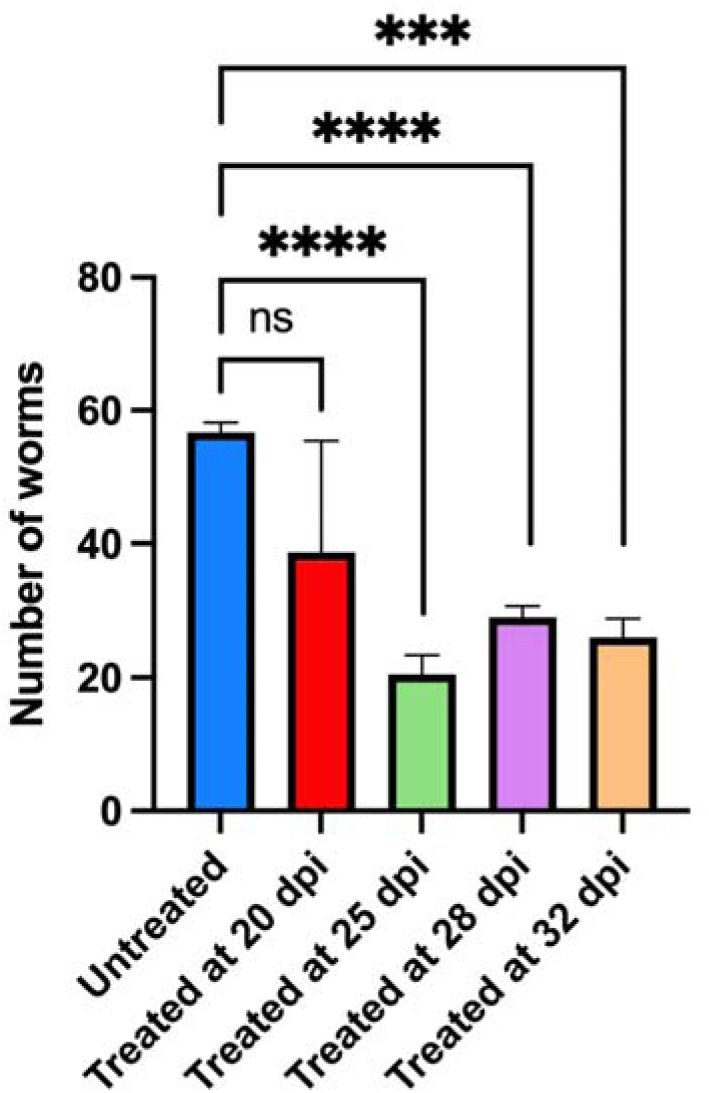
OXA Derivatives Significantly Reduced the Number of Collected *S. mansoni* Juvenile Worms from Infected Mice. Five mice per group were infected with 100 cercaria of *S. mansoni*. Mice were treated with a single dose of 100 mg/kg by oral gavage of CIDD-0150**<u>303</u>** at the day specified on the X-axis and worm burden determined on day 45 pi. Prism/unpaired t test *(P<0.05)*.

The molecular structures of *Sm*SULT with **303** and **610** were determined using X-ray crystallography to characterize their modes of binding in the active site (Fig 7). *Sm*SULT was pre-incubated with the CIDD compounds for 30 min prior to addition of PAP which resulted in crystal complexes of enzyme with bound compounds. The phenyl ring containing the nitro- and hydroxymethyl groups are observed in alternate positions when comparing the compounds to each other and this has been observed previously, likely due to the crystals containing the depleted co-substrate Adenosine 3,5 diphosphate (PAP) instead of the active co-substrate 3 - phosphoadenosine 5′-phosphosulfate (PAPS) which would turn over the substrates [27]. Importantly, the structures revealed that while both **303** and **610** contain indole and (trifluormethyl) phenyl moieties, these groups are interchangeable in position in the substrate binding pocket (Fig 7). These branched moieties occupy two distinct regions in the SULT active site that were observed with previous generations of compounds from our studies that contained single branches [27] which led us to design hybrid/branched generation of compounds.

**Fig 7:**
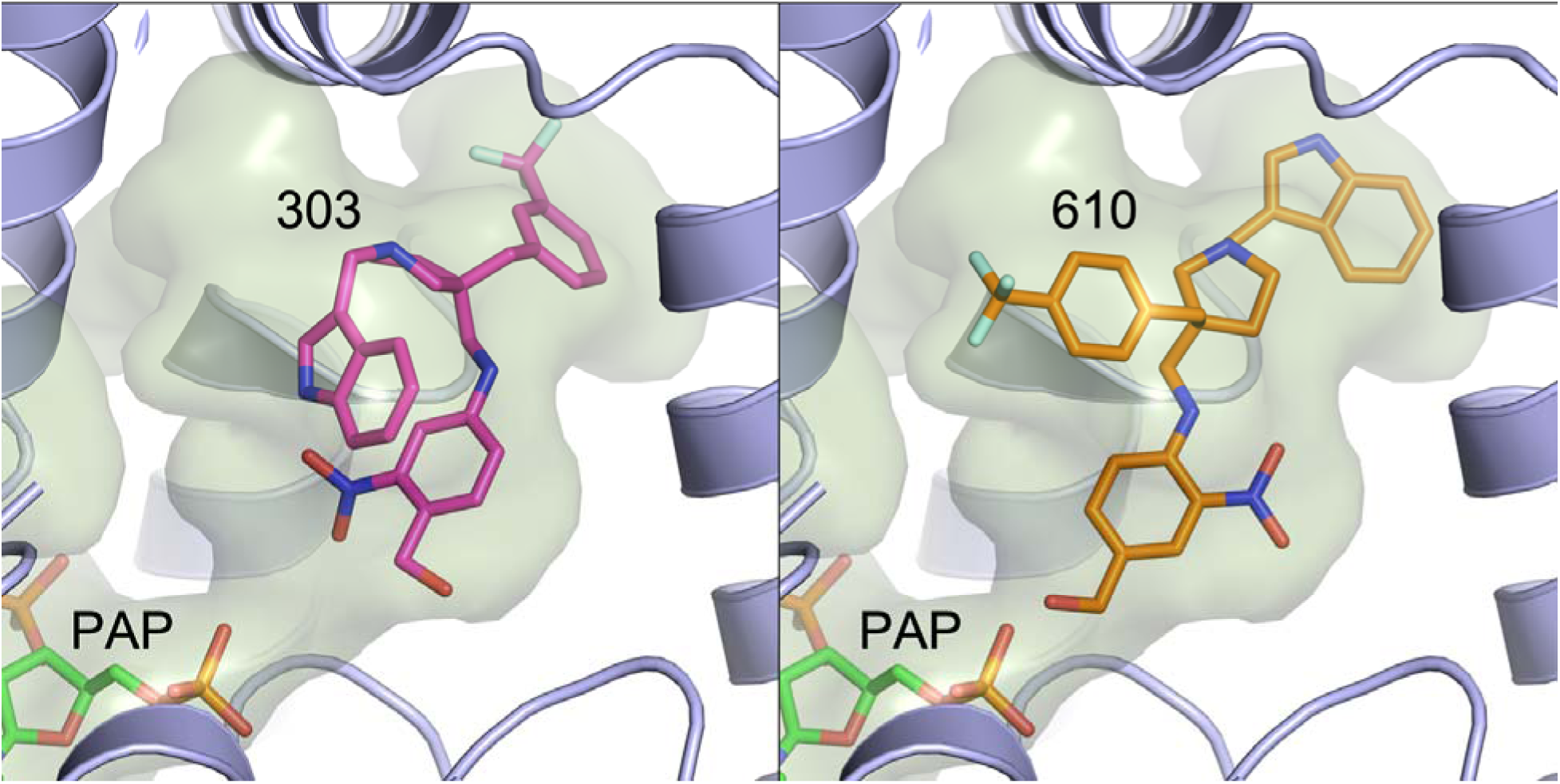
Crystal Structures of CIDD-0150303 (Left Panel) And CIDD-0150610 (Right Panel) Shown in The Active Site of *Sm*SULT. The active site inner surface cavity is depicted in light green. Secondary structure elements in front of the compounds were omitted for clarity. The branched indole and (trifluoromethyl) phenyl groups at top are observed in swapped positions between the two compounds.

To further demonstrate the mechanism of action is through sulfation by a sulfotransferase, we used RNA interference (RNAi) against the *Schistosoma* sulfotransferases (*SmSULT*) and tested the ability of these derivatives to kill *S. mansoni* parasites. The parasites where SULT was knocked down were resistant to drug treatment compared to controls, confirming that the mode of action is conserved [28] (S2 Fig).

PZQ resistance is the most compelling reason for the development of an additional treatment. We have selected PZQ resistant parasites and currently have a PZQ-R isolate that is resistant. IC_50_ for the PZQ-R is 377-fold higher than for the sensitive parasite from which it was derived [31, 32]. We tested OXA derivatives against PZQ-R at a final concentration of 143 µM, **830**, **610**, and **303** were able to kill 100% PZQ-R *in vitro* (Fig 8).

**Fig 8:**
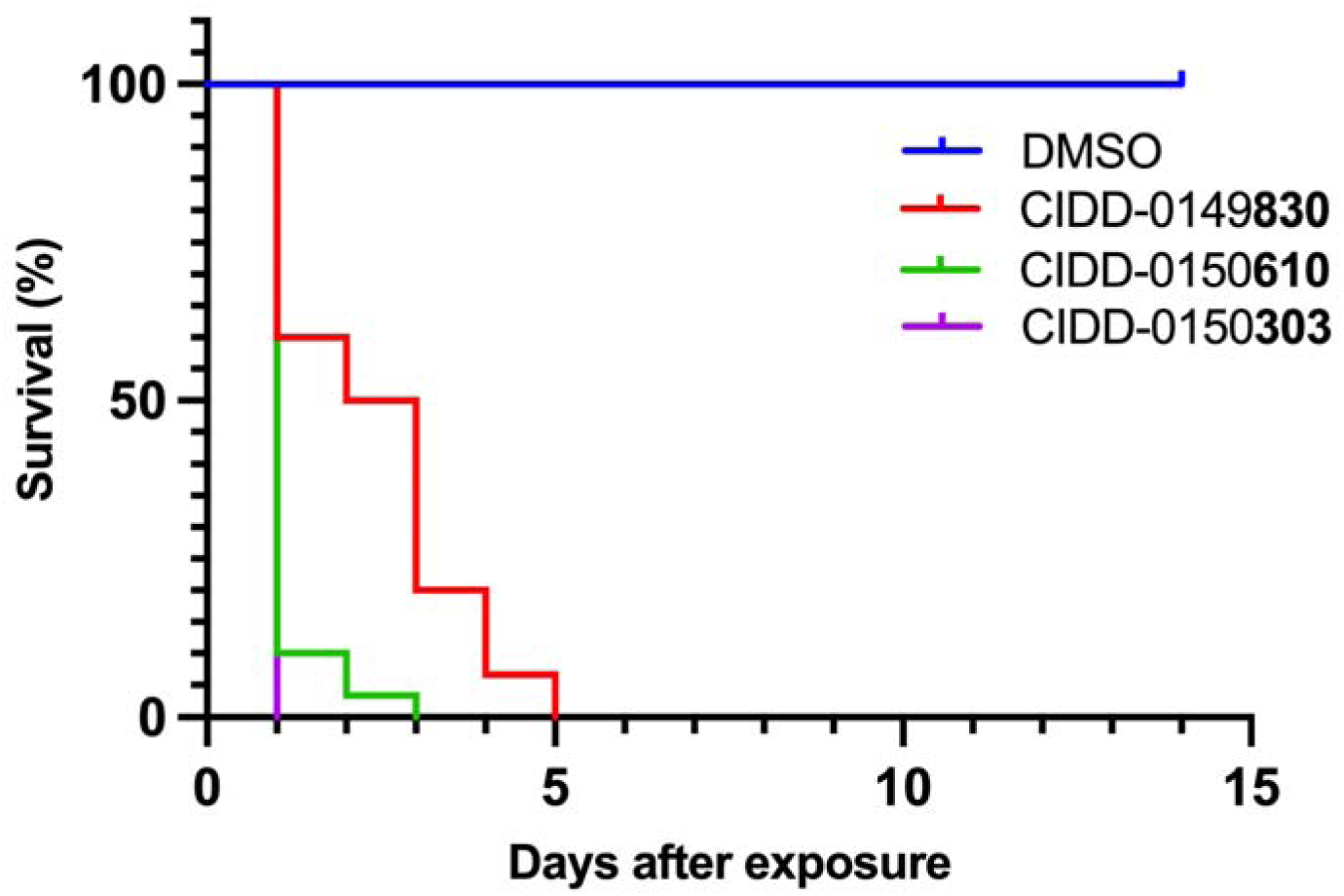
Kaplan-Meier Curves Demonstrate the Ability of CIDD-0149<u>830</u>, CIDD-0150<u>610</u>, and CIDD-0150<u>303</u> to Kill *S. mansoni* PZQ-R. All derivatives were solubilized in 100% DMSO and administered at a final concentration of 143 µM per well for 45 minutes, washed 3 times with media. 45-minute exposure mimics the exposure time in a human Each well contained 10 male adult worms. All screens were performed in experimental and biological triplicate. Survival was plotted as a percentage over time using Prism/Curve Comparison/ Long-rank (Mantel-cox) test. The p-value threshold for each derivative compared to DMSO was <0.001.

To test combination treatment of PZQ and OXA derivatives and demonstrate that the derivatives would kill PZQ-R in an animal model, five mice per group were infected with *S. mansoni* PZQ-R then treated with PZQ, **610**, PZQ + **610**, **303** and PZQ + **303** at a 100 mg/kg of each by oral gavage. Our data shows that the OXA derivative **303** kills PZQ-R worms and that combination treatment of PZQ + **303** significantly reduced the worm burden by 90.8% *P= 0.0001* (Fig 9). The data also demonstrate that the PZQ-R strain was indeed resistant to PZQ as treatment with PZQ did not lead to significant reduction in worm burden.

**Fig 9:**
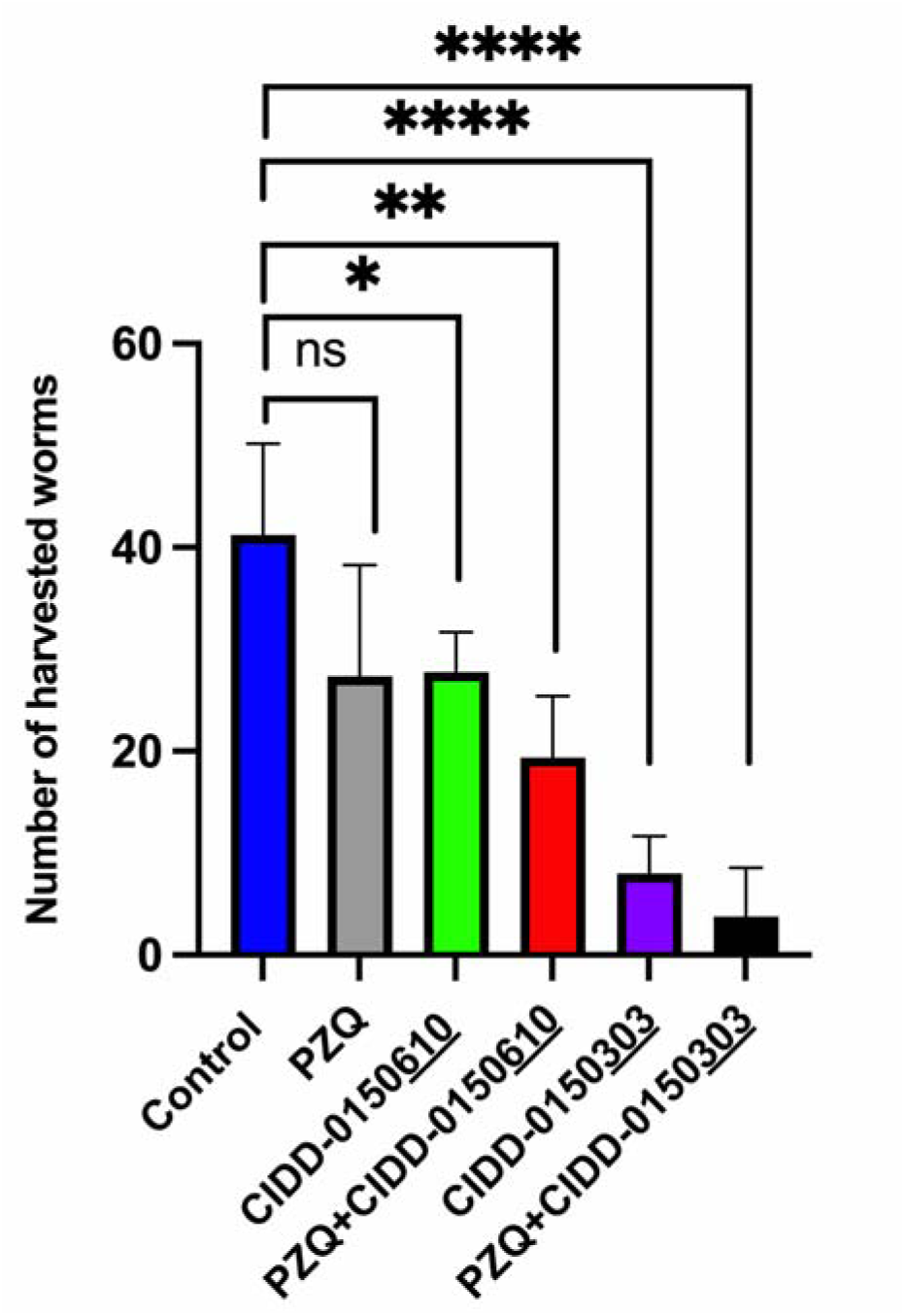
Combination Therapy of PZQ-R Infected Mice. Five mice per group were infected with *S. mansoni* PZQ-R. Worms were collected 10 after days of treatment with a single dose of 100 µg/g by oral gavage and compared to the control group. Prism/unpaired t test *(P<0.05)*.

## Discussion

Our structure-based drug design approach produced a robust Structure Activity Relationship (SAR) program that identified several new lead compounds with effective worm killing [24, 25, 27, 28]. The best derivatives were soaked into *Sm*SULT to repeat this process. This has led to synthesis and testing of more than 350 derivatives of OXA. Of the 350 derivatives, three were identified that kill 100% of all three human schistosomes *in vitro*: **830** [28], **610** and **303** (Fig 1). Moreover, the newly designed OXA compounds were more effective in killing *S. mansoni* than OXA. All the identified compounds kill 100% of *S. mansoni* most requiring less than 14 days, while OXA will kill 90% of *S. mansoni* in 14 days. We chose 143 µM for the *in vitro* screening studies from the calculation of the molarity of 40 mgs/kg, a dose given to humans to be 143 µM. The observation that patient OXA plasma levels are insufficient to kill *S. mansoni in vitro* led to the hypothesis that the OXA levels within the vasculature the worms reside in are more predictive than systemic levels. To support this hypothesis, we performed pharmacokinetic/pharmacodynamic (PK/PD) study for OXA and employing dosing conditions in mice that were modeled and experimentally verified to recapitulate drug exposure observed in human patients treated with OXA, Toth et al., demonstrated that the calculated portal concentration where the schistosomes reside was consistent with the concentrations used in the *in vitro* killing experiment [33]. In addition, the 45 min exposure time used for the *in vitro* experiment mimics the human situation where OXA levels rise immediately after dosing and decline as additional drug is not absorbed from the intestine and systemic OXA is metabolized and excreted [33]. We are working to enhance solubility and bioavailability as we posit that rapid absorption is important for high portal concentration. We previously demonstrated that the difference in sulfotransferases molecular structures among the three *Schistosoma* species does not abrogate OXA binding in the active site. Furthermore, we showed that all three schistosomal enzymes are able to bind to and sulfate OXA to varying degrees *in vitro* [24, 30]. This is linked to the ability of OXA to fit in the binding pocket and to the catalytic efficiency of sulfur transfer [24, 26]. The evolution and progress of OXA derivative design allow for new binding modes for derivatives capable of being active against all three *Schistosoma* species. The efficient and productive binding led to a reduction in the amount of drug required to achieve successful killing (Fig 2). These data show using a dose of 71.5 µM of **610** and **303** was sufficient to kill 100% of *S. mansoni*, *S. haematobium,* and *S. japonicum*. **830** was less effective against *S. haematobium* and *S. japonicum* at 71.5 µM compared to OXA which kills 40% of *S. mansoni* at this concentration. Male worms are 5X more sensitive to OXA than are adult female worms which correlates with the higher levels of *Sm*SULT expression in male adult worms compared to female adult worms [26, 28, 32] However, **303** demonstrates very effective killing for both genders in a very short period as it was able to kill 100% of single mature female and male worms and paired female and male worms all within 24 hours (Fig 3). However, single mature females were less sensitive to **830** and OXA.

We performed an *in vivo* study to evaluate the efficacy of **830**, **610**, **303** in all 3 human schistosome species and **790** in *S. japonicum* (Fig 4). The highest reduction rate among OXA derivatives in *S. mansoni* infection treatment was achieved by **303**. **303** reduced the number of harvested worms by 81.8% *(P=0.017)* compared to the negative control (Fig 4A). All three drugs showed a significant reduction in the number of *S. haematobium* harvested worms. **830** showed a very significant reduction rate 80.2% *(P=0.001)* compared to the negative control. Furthermore, **610** and **303** showed significant killing for *S. haematobium* infection at 69.1% and 60%, respectively (Fig 4B). An effective killing for *S. japonicum* was obtained by **790** and **610** where the reduction rates were 86.7% *(P=0.0003)* and 61% *(P=0.0019)* respectively (Fig 4C). These results are very encouraging as re-engineered OXA is effective against *S. haematobium* and *S. japonicum.* Improving the formulation to enhance aqueous solubility, extend release and prolong uptake will enhance OXA-derivative treatment. Since *S. mansoni* and *S. haematobium* do not occur in China and the Philippines where *S. japoncum* is present, our data suggest that **790** would be the best partner for PZQ to treat *S. japonicum* [33]

Schistosomiasis treatment has a limitation regarding immature worms as PZQ does not kill immature schistosomes allowing the infection to reestablish itself rather quickly [29]. Moreover, previous studies demonstrated a stage-specific susceptibility of *S. mansoni* to OXA treatment [34, 35]. This might be associated with the level of *Sm*SULT transcript. The expression of *Sm*SULT in male worms increases in 21 dpi, 28 dpi reaching the highest levels at day 35 dpi [36–38]. Similar to male worms, female worms’ *Sm*SULT transcript increases in 21 dpi, peaks at day 28 dpi, but then begins to decrease [36, 37]. Therefore, developing a drug that will kill the liver stage schistosomes adds additional value to *Schistosoma* treatment. The effective OXA derivatives; **830** [28], **610** and **303** (Fig 1) demonstrate 100% killing of the liver stage 20-28 dpi worms *in vitro* in 2-6 days. Again, **303** was highly efficacious by killing 100% of liver stages in 1-2 days, *in vitro* (Fig 5). These results were encouraging for evaluating **303** performance in animals (Fig 6). The result, a reduction of worm burden by 49-64%, is an advance over PZQ as PZQ’s lack of efficacy against juvenile schistosomes results in rapid re-infection in highly endemic areas [39]. However, the 20 dpi result was not significant. One possible answer is that immature schistosomes leave the circulation of the lungs and move to the portal circulation of the liver by crossing the splenic bed at around day 18 post infection. Development in schistosomes is asynchronous so some 20 day old worms may not be developed enough to produce sufficient SmSULT to be killed. The X-ray crystal structures of **303** and **610** further demonstrate that the SULT active site can accommodate further modifications in the derivatives. For example, we synthesized two compounds, **303** and **610** with branched indole and (trifluoromethyl)phenyl moieties that swap positions between their binding modes (Fig 7). Obtaining these crystals required pre-incubating apo *Sm*SULT with CIDD compounds prior to adding PAP, which suggests that PAP stabilizes the enzyme structure thereby limiting access to the active site for our larger, branched compounds. However, upon binding, the compounds do not appear to alter the protein structure significantly in the active site or overall. The root-mean-square deviation for the **303** complex compared to OXA-bound *Sm*SULT [40] is 0.50 Å over 1665 atoms and for the **610** complex, 0.16 Å over 1704 atoms calculated using PyMOL. Thus, we anticipate the SULT active site will accommodate further modifications to our most active compounds as we fine tune them to increase favorable pharmacokinetic properties. Our studies demonstrated that OXA derivative **790** has the same mode of action as OXA [38]. Knockdown of *Sm*SULT using RNA-interference (S2 Fig) results in resistance to **830**, **610** and **303** treatments, confirming that the mode of action is conserved. This experiment was a confirmation of our previous work [22] that demonstrates the mode of action of OXA and OXA derivates is the same, due to limited number of worms (n=30), we performed qPCR only once therefore we didn’t show the figures. In contrast to OXA, the PZQ mode of action was not completely understood [41], until recently [31, 32]. It is now known that PZQ activates a flatworm transient receptor potential channel (TRPMPZQ) to mediate sustained Ca2+ influx and worm paralysis [32]. Thus, PZQ mode of action is different than OXA derivatives. We did an experiment with a susceptible strain and treated with 100 mg/kg PZQ. We obtain about 90% killing. Treated PZQ-resistant parasites (PZQ-R) with **830** [28], **610** and **303** results in 100% killing *in vitro* (Fig 8). These results encouraged the evaluation of PZQ and the best OXA derivates in combination to treat PZQ-R infected mice. Combination therapy of PZQ + **303** resulted in 90.8% reduction in the PZQ-R worm burden (*P= 0.0001,* Fig 9) which strengthens our drug discovery outcomes since PZQ and **303** have a different mode of action. Schistosomes are dioecious multicellular eucaryotic parasites that do not multiply within the human body but reproduce sexually producing eggs which are responsible for pathogenesis and transmission. Drug resistance to PZQ and OXA are double recessive traits [31, 42]. The chance that an adult male and separately an adult female would develop resistance to both drugs is remote.

We conclude that **830**, **790** and **303** are potential new drugs to treat *S. haematobium* and *S. japonicum.* **303** is the potential drug that can be used with PZQ in combination for better treatment and to mitigate the development of resistance. The research now focuses on physicochemical, ADME, pharmacokinetics and toxicology studies that will justify requesting authorization from the Food and Drug Administration and ultimately clinical trials.

## Materials and Methods

### Parasite Maintenance

*Schistosoma mansoni*, *S. haematobium* and *S. japonicum* were maintained by passage through a snail intermediate host, *Biomphalaria glabrata*, *Bulinus truncatus* or *Oncomelania hupensis*, respectively. Golden Syrian Hamsters were the definitive host. Hamsters were infected with 250 cercariae of *S. mansoni, S. haematobium, or S. japonicum.* to maintain the schistosome life cycle of each species according to IACUC protocol (Protocol #08039).

### Parasite Recovery

Depending on *Schistosoma* species and the required stages for each experiment, the infected hamsters were sacrificed between 30 to 90 days post-infection (dpi) in accordance with IACUC protocol (UTHSCSA IACUC Protocol #08039). S. japonicum at 30 days, S. mansoni at 45 days and S. haematobium at 90 days. Animals were euthanized by intraperitoneal injection using Fatal-Plus (Butler Animal Health, Ohio), a sodium pentobarbital solution, and 10% heparin. Adult schistosomes were collected by perfusion as previously described (Duvall et al., 1967) using 0.9% saline containing EDTA.

### Parasite *In Vitro* Culture

Harvested worms were cultured in 2ml 1X Dulbecco’s Modified Eagle Medium (DMEM, Gibco) with 10% Heat Inactivated Fetal Bovine Serum (FBS, Atlantic Biologicals) and 1X antibiotic/antimycotic (Ab/Am, GIBCO). Worms were manually sorted under a dissecting stereomicroscope and aliquoted to 10 single worms or paired worms per well in a 24-well plate. Three-hour schistosomula were mechanically transformed from cercariae according to Tucker et al. [43]. Worms were cultured in an incubator at 37°C and 5% CO_2_ for 72 hours. Worm viability was assessed by daily observation. Culture media was changed every other day.

### OXA Derivative Design and Synthesis

We employed an iterative process to develop new drugs [25]. To do this **830** was soaked into *Sm*SULT crystals, the resulting SARs information was used by the Center for Innovative Drug Discovery (CIDD) to synthesize new derivatives that were tested for schistosomicidal activity in an *in vitro* killing assay [25, 27, 28]. The synthesis of the **830** chemical series was previously published [25].

### OXA Derivative *In Vitro* Assays

OXA derivatives were solubilized in 100% Dimethyl sulfoxide (DMSO) then diluted to reach to the final concentrations 14.3 µM, 35.75 µM, 71.5 µM and 143 µM depending on the experiment’s purposes. The derivatives were added directly to each well within 2-4 hours after collecting schistosomes from the hamsters. Each derivative was tested in triplicate. In addition to evaluate the derivatives efficacy at 143 µM and determine the minimum dose, we tested the ability of OXA derivatives to kill both genders and to kill Juvenile worms. Harvested worms from bisex infection were sorted to single female and male worms and female and male worms in worm pairs. To evaluate the ability of OXA derivatives to kill Juvenile stages male worms were collected in 20 dpi, 25 dpi, 28 dpi, 32 dpi, 45 dpi. Worms collected at 20 dpi were not sorted by sex. OXA was the positive control for only *S. mansoni* [18]. Drugs were incubated with schistosomes at 37°C, 5% CO2 for 45 minutes, mimicking physiological conditions [22]. The worms were washed with plain media 3 times to remove any residual derivatives. Worms were then incubated in culture media for a period of up to 14 days. Worm motility, tegument shedding, opaque color, and tegument blebbing were used to evaluate survival and death/morbidity. Worms were observed daily up to 14 days. They were considered dead when they showed a lack of motility especially no response to being poked and were opaque. Culture media was changed every other day.

### OXA Derivative *In Vivo* Assays

#### Evaluate the ability of OXA derivatives to kill the main three *Schistosoma* species

Five Balb/c mice per group were infected with 80 cercaria *S. mansoni* and maintained for 45 dpi, then treated by gavage with single oral dose of 100 mg/kg of OXA derivatives dissolved in 5% DMSO and 95% ethanol. Animals were perfused 10 days after treatment and the number of harvested worms counted. Control groups were treated with either diluent or OXA. To evaluate the ability of OXA derivatives to kill *S. haematobium* and *S. japonicum* in animal models 5 hamsters per group were infected with 100 cercariae and maintained for 90 dpi and 30 dpi, respectively. Then animals were treated by gavage with single oral dose of 100 mg/kg of OXA derivative dissolved in 5% DMSO and 95% ethanol. The control groups were treated with diluent. Ten days after treatment, animals were perfused and collected worms were counted and compared to the control.

#### Evaluate 303 to kill immature, liver stage schistosomes

Five Balb/c mice per group were infected with 100 cercaria of *S. mansoni*. Mice (n=5) were treated on days 20 dpi, 25 dpi, 28 dpi or 32 dpi with 100 mg/kg **303** and perfused on day 45 pi. The number of harvested worms from each treatment group were compared to untreated mice. **303** was dissolved in 5% DMSO and 95% Ethanol.

#### Test combination treatment of PZQ and OXA derivatives ability to kill *S. mansoni* PZQ-R

To test PZQ in combination with OXA derivatives five Balb/c mice per group were infected with 80 cercaria of *S. mansoni* PZQ-R. Mice were treated with 100 mg/kg by oral gavage with single oral dose of PZQ, **610**, **303**. For combination treatment groups mice were treated with single oral dose of PZQ + **610** and PZQ + **303** with 100 mg/kg of each drug. Drugs were dissolved in 5% DMSO and 95% ethanol. The animals were treated at 45 dpi and *S. mansoni* PZQ-R worms were harvested at day 55 post infection. The control groups were treated with diluent.

### RNA Extraction

Total RNA was obtained from frozen samples of adult *S. mansoni* worms. All frozen samples were thawed on ice in RNAzolRT (Molecular Research Center Inc.) each sample then was placed in 2 ml tubes of Lysin Matrix Tubes containing 1.4 mm ceramic spheres and then homogenized 2x using Beadbeater homogenizer (Biospec, USA) for 45 seconds. RNA was extracted and purified according to (Molecular Research Center Inc.) manufacturer instructions for total RNA isolation.

### cDNA Synthesis

cDNA was generated from total RNA using BioRad iSCRIPT cDNA Synthesis Kit according to the manufacturer’s instructions.

### dsRNA Synthesis and Treatment

Forward 5’-ATT GGA TGG TTA CAT AGC AAC TAC-3’ and reverse 5’-CCA TGG ATC ATT TGA TTT GGG T-3’ primers amplifying a 192-592 bp section of the coding region for *Sm*SULT (Smp_ 089320) were designed using PrimerDesign tool by IDTdna. Polymerase chain reaction (PCR) was performed to produce an amplicon, followed by confirmation of amplification by running the PCR product on a 1% agarose gel.

T7 promoters were added to the forward and reverse primer to flank the PCR product. Confirmation of amplification was also performed via 1% agarose gel. The PCR product with T7 promoters were used as a template for transcription of the dsRNA. The dsRNA was placed in a 37°C water bath within 24 hours and treated with DNAase to remove contaminants. Ammonium acetate 3 M was added, followed by 100% ethanol to precipitate the RNA. The RNA was left at this step overnight. Then the sample was centrifuged at 14000 rpm, forming an RNA pellet. The pellet was washed twice with 70% ethanol. On the second wash, the supernatant was removed, and the ethanol allowed to evaporate. Then the pellet was resuspended in nuclease-free water. The concentration of RNA was then measured using the Thermo Scientific NanoDroprop 1000 spectrophotometer.

Ten adult male schistosomes were collected, sorted, and treated at 45 dpi with 30 µg/mL dsRNA in triplicate of *S. mansoni* SULT or irrelevant control *Luciferase* (M15077) right after worm sorting. Then worms were treated again after 3, 7, and 11 days. OXA derivatives (143 µM) were added at day 6. The worms were observed for 14 days. Observation included notes on worm health, viability; lack of motility, shedding of tegument, blebbing of tegument, internal vacuolization, lethargy, and being opaque [38].

### Crystallization, Structure Determination, and Refinement of CIDD-0150610 and CIDD-0150303 Complexed with SULT

Automated screening for crystallization was carried out using the sitting drop vapor-diffusion method with an Art Robbins Instruments Phoenix system in the Structural Biology Core at the University of Texas Health Science Center at San Antonio. *Sm*SULT was prepared as previously described [22]. CIDD compounds were added first to apo *Sm*SULT and incubated for 30 min prior to adding PAP. The protein complexes were mixed in a 1:1 ratio with crystal screen reagents for a total drop volume of 0.4 mL. *Sm*SULT:CIDD-0150**303** crystals were grown at 22°C in Molecular Dimensions Morpheus condition 1-1 (30% Precipitant Mix 1 [30% PEG 500 MME; 20% PEG 20000], 0.06 M Divalents Mix [0.3 M magnesium chloride hexahydrate, 0.3 M calcium chloride dihydrate], 0.1 M Buffer System 1 pH 6.5 [1.0 M imidazole:MES]). *Sm*SULT:CIDD-0150**610** crystals were grown at 4°C in Anatrace MCSG-3 condition A1 (20% PEG 8000, 0.1 M HEPES:NaOH pH 7.5). Crystals were flash-cooled in liquid nitrogen by wicking off excess solution from crystals harvested in nylon cryo-loops prior to data collection at the Advanced Photon Source, Argonne, IL, NE-CAT beamline 24-ID-E. Diffraction data were processed using AUTOPROC [44]. The structures were determined by the molecular replacement method implemented in PHASER[45] using coordinates from PDB entry 6BDR [27]as the search model. Coordinates were refined using PHENIX [46] including simulated annealing and alternated with manual rebuilding using COOT [47]. All models were verified using composite omit map analysis [48]. Data collection and refinement statistics are shown in Table S2. PyMOL was used to generate images for the crystal structures (The PyMOL Molecular Graphics System, Version 2.2 Schrödinger, LLC.).

### Statistical Analysis

Statistical analysis for the Kaplan-Meier curves were performed using GraphPad Prism software (version 9.3.1). Differences in the survival function of different treatments were tested using a Curve Comparison/ Long-rank (Mantel-cox) test. Unpaired t test was used for treatment comparisons in animal models.

## Acknowledgments

*Oncomelania hupensis* exposed to *S. japonicum* were provided by the Schistosomiasis Resource Center of the Biomedical Research Institute (Rockville, MD) through NIH-NIAID Contract HHSN272201700014I:. We thank the Structural Biology Core Facilities, a part of the Institutional Research Cores at the UT Health at San Antonio and Advanced Photon Source, a U.S. Department of Energy (DOE) Office of Science User Facility operated for the DOE Office of Science by Argonne National Laboratory for using their resources. We also acknowledge Ovidiu Dumitru Orbai for assistance with experimental procedures.

## Funding

San Antonio Biomedical Education and Research Program. Institutional Research and Academic Career Development Award from the National Institute of General Medical Science 2k12GM111726 Postdoctoral support for SA.

National Institute of Allergy and Infectious Diseases AI27219 to PTL.

Morrison Trust to PTL.

Institutional Research Cores at the University of Texas Health Science Center at San Antonio supported by the Office of the Vice President for Research, Greehey\: Children’s Cancer Research Institute and Mays Cancer Center Drug Discovery and Structural Biology Shared Resource National Institutes of Health grant P30 CA054174 to ABT. The Rigaku HyPix-6000HE Detector, Universal Goniometer and VariMax-VHF Optic instrumentation in the Structural Biology Core Facilities are funded by NIH-ORIP SIG Grant S10OD030374 to ABT.

National Institute of General Medical Sciences from the National Institutes of Health (P30 GM124165) to ABT.

The Eiger 16M detector on the 24-ID-E beam line is funded by a NIH-ORIP HEI grant (S10OD021527) to ABT.

## Supporting Information

**S1 Fig. Ability of OXA And OXA Derivatives to Kill *Schistosoma* Species at Final Concentrations of 143 µm, 71.5 µm, 35.75 µm, And 14.3 µm Per Well *In Vitro*.** A. OXA against *S. mansoni*. B.1. CIDD-0149**830** against *S. mansoni*, B.2. CIDD-0149**830** against *S. haematobium.* B.3. CIDD-0149**830** against *S. japonicum.* C.1. CIDD-0150**610** against *S. mansoni*, C.2. CIDD-0150**610** against *S. haematobium* and C.3. CIDD-0150**610** against *S. japonicum.* D.1. CIDD-0150**303** against *S. mansoni*, D.2. CIDD-0150**303** against *S. haematobium, and* D.3. CIDD-0150**303** against *S. japonicum.* E. The percentage of worms killed at each concentration. OXA and OXA derivatives were tested against adult male worms. All drugs were solubilized in 100% DMSO. All screens were performed in experimental and biological triplicate. Survival was plotted as a percentage over time using Prism/Curve Comparison/ Long-rank (Mantel-cox) test. The p-value threshold for each derivative compared to DMSO was <0.001

**S2 Fig. Kaplan-Meier Curves Demonstrate the Knockdown of *S. mansoni* Sulfotransferase Confers Resistance Upon Challenge.** A.1. CIDD-0149<u>830</u>: *Sm*SULT RNAi alone, Irrelevant RNAi, *Sm*SULT RNAi + OXA, and *Sm*SULT RNAi + **830** had 93%+ survival and were displaying healthy characteristics. All other groups expressed similar, expected sensitivity levels to **830** treatments A.2. CIDD-0150**<u>610</u>**: *Sm*SULT RNAi alone, Irrelevant RNAi, and *Sm*SULT RNAi + **610** had 90%+ survival and were displaying healthy characteristics. All other groups expressed similar, expected sensitivity levels to **610** treatments A.3. CIDD-0150<u>303</u>: *Sm*SULT RNAi alone, Irrelevant RNAi, and *Sm*SULT RNAi + **303** had 93%+ survival and were displaying healthy characteristics. All other groups expressed similar, expected sensitivity levels to **303** treatments.

**S1 Table. Chemical Structure Of CIDD-066<u>790,</u> CIDD-0149<u>830,</u> CIDD-0150<u>610</u>, and CIDD-0150<u>303</u>.**

**S2 Table. Crystallographic Data Collection and Refinement Statistics.**

**S3 Table. OXA Derivatives Against Schistosoma Species In Vitro Results**

**S4 Table. Test The Efficacy of OXA Derivates in An *in Vivo* Model.**

