## Supplemental data for "Oxamniquine Derivatives Overcome Praziquantel Treatment Limitations for Schistosomiasis"

**Supporting Information (for publication)**

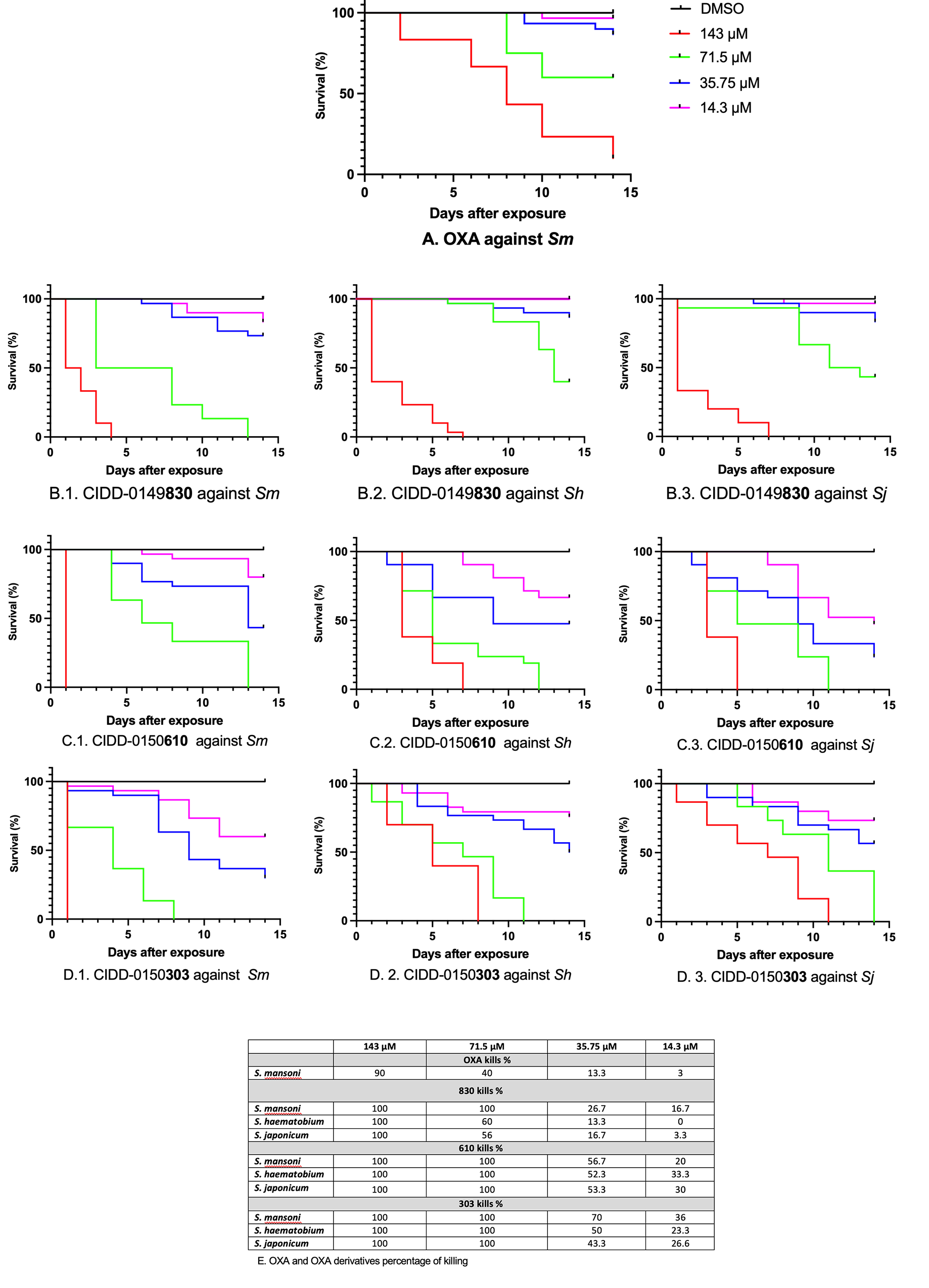

**S1 Fig. Kaplan-Meier Curves Demonstrate the Ability of OXA And OXA Derivatives to Kill *Schistosoma* Species at Final Concentrations of 143 µm, 71.5 µm, 35.75 µm, And 14.3 µm Per Well *In Vitro.*** A. OXA against *S. mansoni*. B.1. CIDD-0149830 against *S. mansoni*, B.2. CIDD-0149830 against *S. haematobium.* B.3. CIDD-0149830 against *S. japonicum.* C.1. CIDD-0150610 against *S. mansoni*, C.2. CIDD-0150610 against *S. haematobium* and C.3. CIDD-0150610 against *S. japonicum.* D.1. CIDD-0150303 against *S. mansoni*, D.2. CIDD-0150303 against *S. haematobium, and* D.3. CIDD-0150303 against *S. japonicum.* E*.* The percentage of worms killed at each concentration. OXA and OXA derivatives were tested against adult male worms*.* All drugs were solubilized in 100% DMSO. All screens were performed in experimental and biological triplicate. Survival was plotted as a percentage over time using Prism/Curve Comparison/ Long-rank (Mantel-cox) test. The p-value threshold for each derivative compared to DMSO was <0.001.

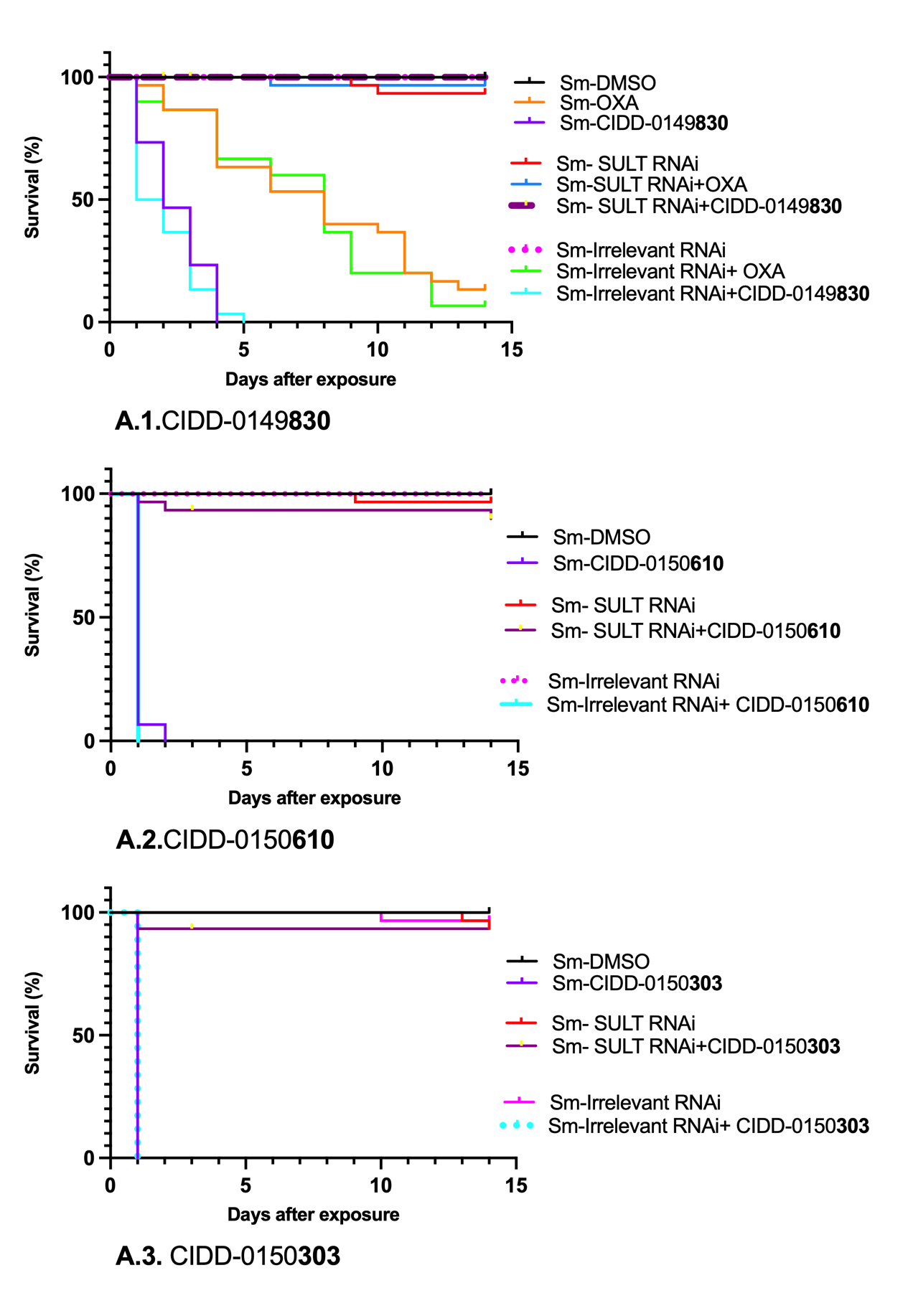

**S2 Fig**. **Kaplan-Meier Curves Demonstrate the Knockdown of *S. Mansoni* Sulfotransferase Confers Resistance Upon Challenge.** A.1. CIDD-0149830**:** *Sm*SULT RNAi alone, Irrelevant RNAi, *Sm*SULT RNAi + OXA, and *Sm*SULT RNAi + 830 had 93%+ survival and were displaying healthy characteristics. All other groups expressed similar, expected sensitivity levels to 830 treatments A.2. CIDD-0150610: *Sm*SULT RNAi alone, Irrelevant RNAi, and *Sm*SULT RNAi + 610 had 90%+ survival and were displaying healthy characteristics. All other groups expressed similar, expected sensitivity levels to 610 treatments A.3. CIDD-0150303**:** *Sm*SULT RNAi alone, Irrelevant RNAi, and *Sm*SULT RNAi + 303 had 93%+ survival and were displaying healthy characteristics. All other groups expressed similar, expected sensitivity levels to 303 treatments.

**S1 Table. Chemical Structure Of CIDD-066790,** **CIDD-0149830, CIDD-0150610, and CIDD-0150303.**

| **OXA derivatives** | **Chemical Structure** |
| --- | --- |
| CIDD-066**790** | 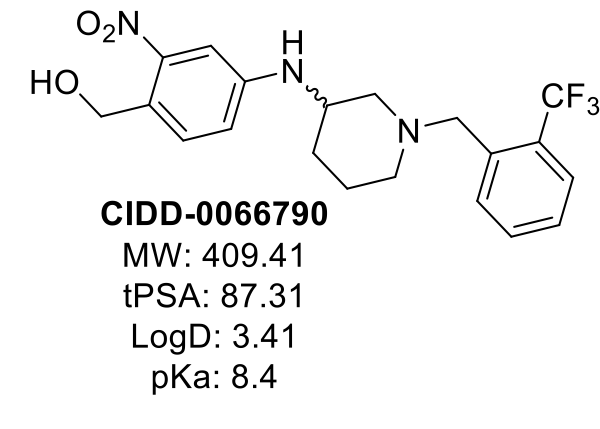 |
| CIDD-0149**830** | 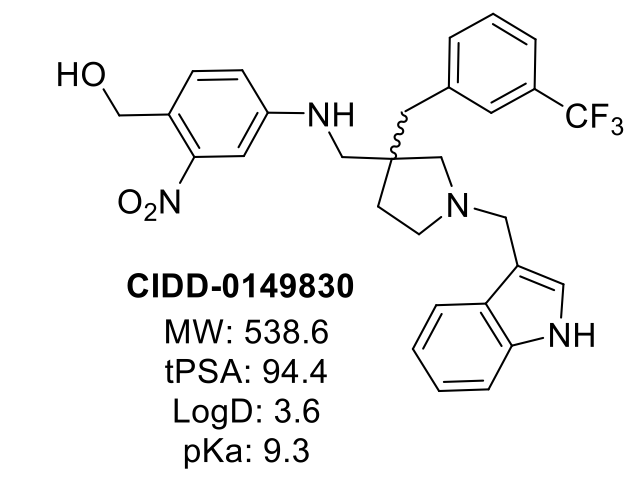 |
| CIDD-0150**610** | 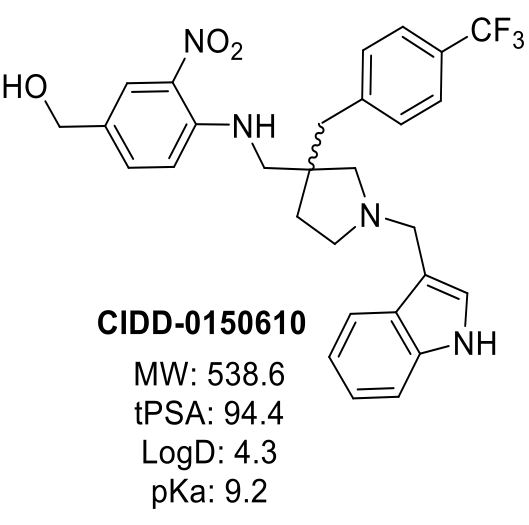 |
| CIDD-0150**303** | 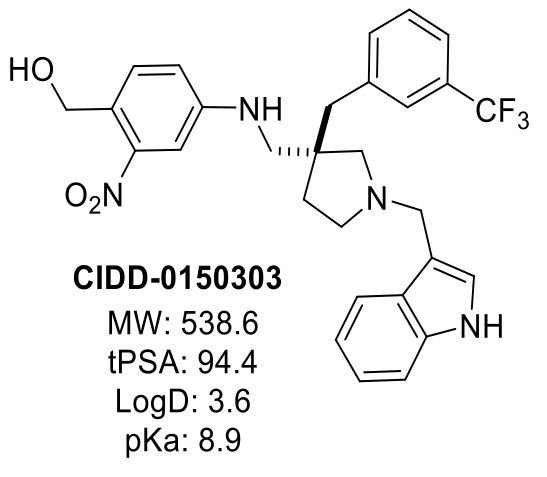 |

**S2 Table.** **Crystallographic Data Collection and Refinement Statistics.**

| **Data collection** |  |  |
| --- | --- | --- |
| PDB Code | 8E5Q | 8E5R |
| Ligands | CIDD-0150303 | CIDD-0150610 |
| Space group | *P*2_1_2_1_2_1_ | *P*2_1_2_1_2 |
| Cell dimensions |  |  |
| *a*, *b*, *c* (Å) | 46.8, 58.5, 90.5 | 140.9, 39.7, 53.8 |
| α, β, γ (°) | 90, 90, 90 | 90, 90, 90 |
| Wavelength (Å) | 0.97918 | 0.97918 |
| Resolution (Å) | 90.53-1.33 (1.40-1.33)* | 140.89-1.40 (1.48-1.40) |
| *R*_pim_ | 0.025 (0.785) | 0.022 (0.817) |
| CC_1/2_ | (0.557) | (0.515) |
| Mean *I* / σ*I* | 13.6 (1.0) | 13.3 (1.0) |
| Completeness (%) | 99.6 (98.7) | 97.6 (99.9) |
| Redundancy | 6.2 (6.3) | 6.0 (6.2) |
| Wilson value (Å^2^) | 18.3 | 23.7 |
| **Refinement** |  |  |
| Resolution (Å) | 49.13-1.33 (1.36-1.33) | 53.84-1.40 (1.44-1.40) |
| No. reflections | 57,487 | 58,966 |
| *R*_work_ / *R*_free_ | 0.182/0.217 | 0.169/0.199 |
| No. atoms |  |  |
| Protein | 2,015 | 2,113 |
| Ligands | 66 | 84 |
| Solvent | 215 | 222 |
| *B*-factors (Å^2^) |  |  |
| Protein | 32.5 | 33.4 |
| Ligand | 34.5 | 41.5 |
| Solvent | 40.0 | 41.6 |
| R.m.s. deviations |  |  |
| Bond lengths (Å) | 0.007 | 0.013 |
| Bond angles (°) | 1.049 | 1.247 |
| Ramachandran Plot |  |  |
| Favored (%) | 96.62 | 96.84 |
| Allowed (%) | 3.38 | 3.16 |
| Outliers (%) | 0.00 | 0.00 |

*Values in parentheses are shown for the highest resolution shell.

**S3 Table. OXA Derivatives Against Schistosoma Species *In Vitro* Results**

|  | **OXA** | **830** | **610** | **303** |
| --- | --- | --- | --- | --- |
| Killing *Schistosoma* species | *S. mansoni* | *S. mansoni*  *S. haematobium*  *S. japonicum* | *S. mansoni*  *S. haematobium*  *S. japonicum* | *S. mansoni*  *S. haematobium*  *S. japonicum* |
| Dose require to kill 100% of *S. mansoni* | 143 µM kills 90% | 71.5 µM | 71.5 µM | 71.5 µM |
| Dose require to kill 100% of *S. haematobium* | Not effective | 143 µM | 71.5 µM | 71.5 µM |
| Dose require to kill 100% of *S. haematobium* | Not effective | 143 µM | 71.5 µM | 71.5 µM |
| Killing adults of single sex- female and male worms and female and male worms in worm pairs of ***S. mansoni*** in 143 µM | Effective killing for both gender except paired males 60% of killing | 100% of killing of both gender except  paired females 60% of killing | 100% of killing of both gender | 100% of killing of both gender |
| Killing adults of single sex- female and male worms and female and male worms in worm pairs of ***S. haematobium*** in 143 µM | Not effective | 100% of killing of both gender except  paired females 63% of killing | 100% of killing of both gender | 100% of killing of both gender |
| Killing adults of single sex- female and male worms and female and male worms in worm pairs of ***S. japonicum*** in 143 µM | Not effective | 100% of killing of both gender except  paired females 70% of killing | 100% of killing of both gender | 100% of killing of both gender |
| Killing juvenile worms of ***S. mansoni*** in 143 µM | Not effective (data not shown) | 100% of killing | 100% of killing | 100% of killing |
| Killing **PZQ-resistant strain** in 143 µM | Not tested | 100% of killing | 100% of killing | 100% of killing |

**S4 Table. Test The Efficacy of OXA Derivates in An *in Vivo* Model.**

| A-The reduction in worm burden after treatment with OXA derivatives against *Schistosoma* species (5 animals were treated with a single dose by oral gavage with 100 mg/kg) | | | | | |
| --- | --- | --- | --- | --- | --- |
|  | **OXA** | **830** | **610** | **303** | **790** |
| *S. mansoni* | 93% | 72.3% | 47% | 81.8% |  |
| *S. haematobium* | Not effective | 80.2% | 69.1% | 60% |  |
| *S. japonicum* | Not effective | 38.3% | 61% | 31% | 86.7 |
| B- The ability of at 100 mg/kg as a single oral dose **303** to kill juvenile worms in an *in vivo* study (5 animals were treated with a single dose by oral gavage with 100 mg/kg) | | | | | |
| Day post infection | | 20 | 25 | 28 | 32 |
| Reduction in worm burden | | NS | 63.8% | 48.9% | 54.1% |
| C- Combination treatment of PZQ and OXA derivatives against PZQ-resistant strain (5 animals were treated with a single dose by oral gavage with 100 mg/kg of each) | | | | | |
| Combination treatment | | PZQ + **610** | | PZQ + **303** | |
| Reduction in worm burden | | 52.9 % | | 90.8% | |
